# Relationship between ACTN3 and ACE polymorphisms and FIFA World averaged rating

**DOI:** 10.1101/363358

**Authors:** Valery Ilinsky, Natalia Zakharova

**Affiliations:** Genotek Ltd.; Pirogov Russian National Research Medical University, Moscow, Russia

## Abstract

Polymorphisms in ACTN3 and ACE genes were previously reported to be associated with power and endurance performance. We estimated the relationship between their population frequencies in several countries and FIFA World averaged rating (FWAR). The entropy was used as the measure of uncertainty. The FWAR distribution entropy (in nats) was 1.38. The FWAR conditional distribution entropy given ACTN3 RR and ACE II frequencies was 1.27 and 1.24, respectively. Almost same FWAR conditional entropy (1.25) was after adjustment on GDP. Population frequencies of polymorphisms in power and endurance performance genes might be a cause of high or low FWAR (low frequency -- less football players) or an effect of ethnic origin and geographical location of countries, related to GDP.

## Introduction

Football involves physical activities of various nature. Professional players cover about 8-12 kilometers during 90-min match [1]. A large amount of this distance corresponds to walking and light running, and up to 20% corresponds to maximal or near maximal running speed during the critical phases of games. Because of the very high intensity of critical game phases, and high level of endurance capacity both power and endurance abilities are critical to achieve success in football. Recent genetic studies of elite athletes have been identified several loci contributing to endurance and power performance [2].

Polymorphisms in α-actinin-3 (ACTN3) and angiotensin I converting enzyme (ACE) genes are associated with power and endurance abilities, respectively [3, 4]. A premature stop codon SNP (p.Arg577Ter, rs1815739) in ACTN3 leads to a lower level of α-actinin-3. The expression of α-actinin-3 is almost exclusively restricted to type II (fast) muscle fibres, therefore alpha-actinin-3 deficiency (XX, rs1815739(T/T) genotype) appears to impair power performance. The frequency of two functional copies of ACTN3 (RR, rs1815739(C/C) genotype) was significantly higher among elite footballers compared the sedentary controls [5].

Angiotensin I converting enzyme encoded by ACE gene catalyzes the conversion of angiotensin I into a physiologically active angiotensin II controlling blood pressure. Indel polymorphism of ACE gene has been shown to be associated with enzyme activity and endurance phenotype [6].

National soccer team success depends on professional level of individual players, but their average level depends on various factors, including popularity of soccer in general, accessibility of soccer schools for young athletes, etc. We estimated the impact of ACTN3 rs1815739 and ACE Indel polymorphisms frequencies as well as GDP on national soccer teams FIFA World averaged rating (FWAR) [7].

## Methods

The ACTN3 rs1815739 and ACE I/D allele frequencies in countries for which FWAR is available were collected using data of scientific papers [8,9] and open databases, such as Alfred [10]. The frequencies of the genetic markers in Russian population were obtained from Genotek database.

Entropy is a common way to describe uncertainty of a random variable. Higher entropy is intrinsic to a more “surprisal” events. Low values of FWAR entropy indicate expected outcome due to the nature of football teams. Entropy of FWAR distribution was calculated using R package “infotheo_1.2.0.” [11].

## Results

The scatter plots of ACTN3 RR (rs1815739(C/C)) and ACE I/I frequencies and FWAR are presented on figure 1 and figure 2 respectively.

**Figure 1.**
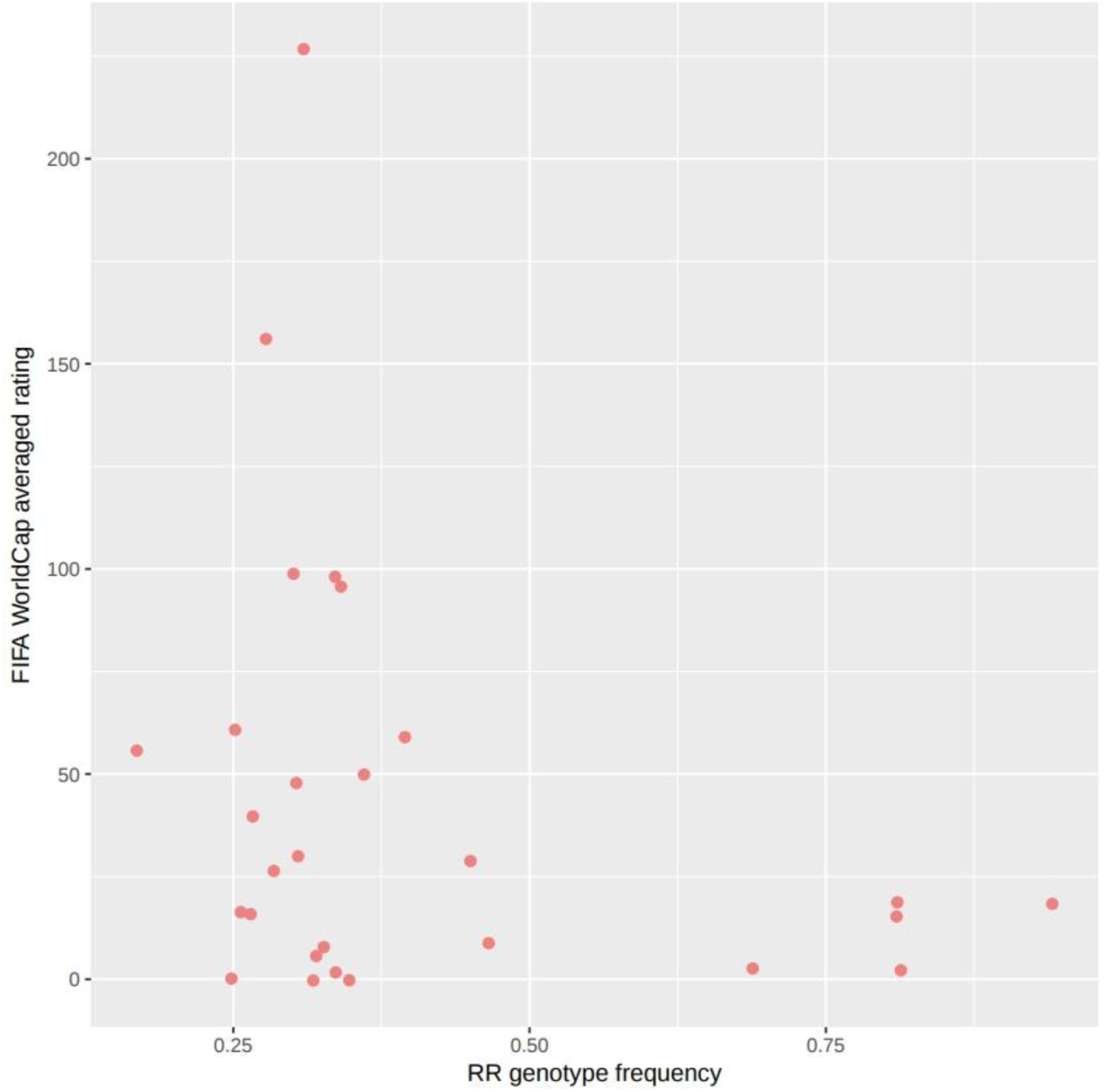
Scatter plot of correlation between ACTN3 RR (rs1815739(C/C)) genotype and FIFA World averaged rating.

**Figure 2.**
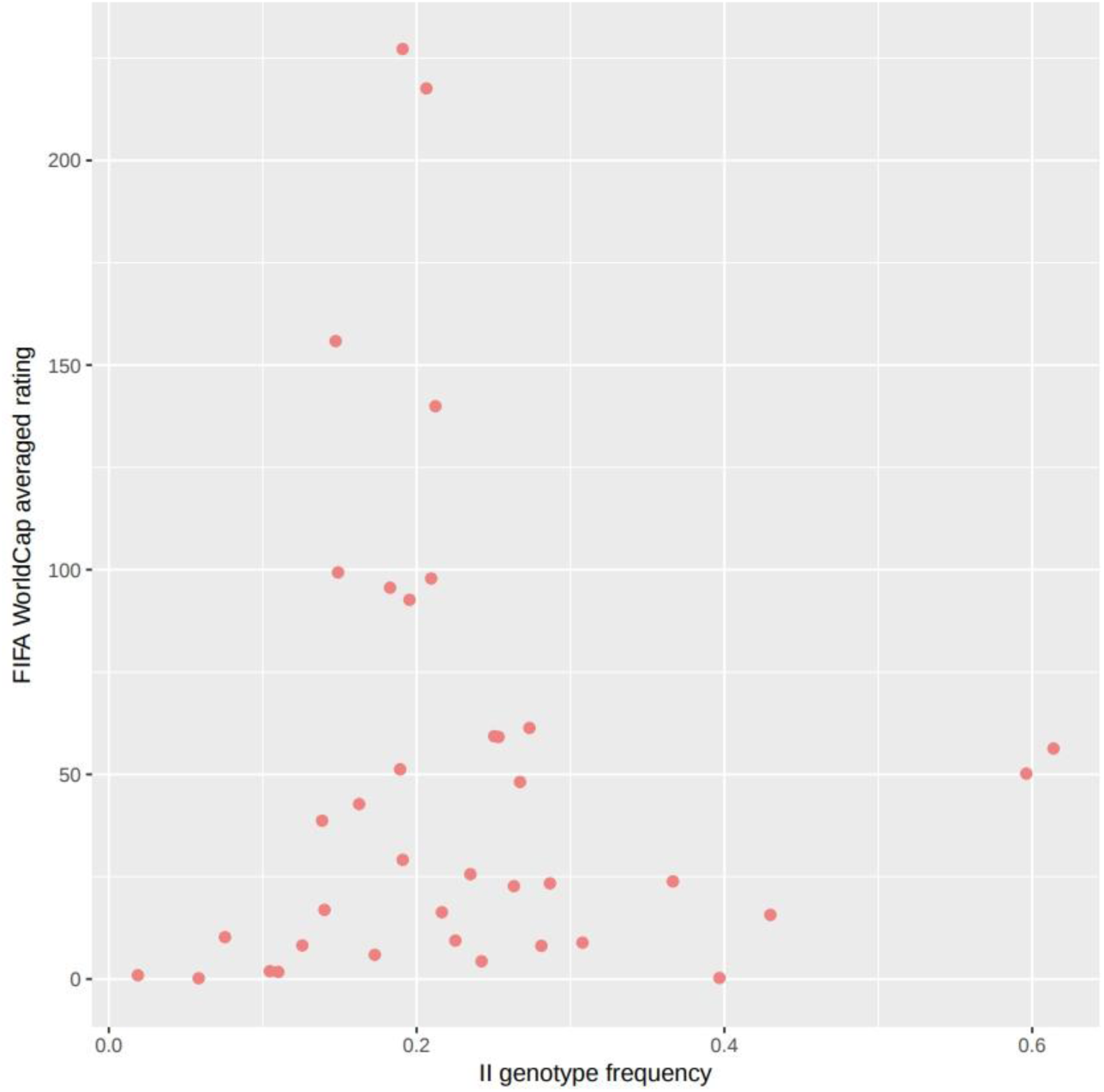
Scatter plot of correlation between ACE II genotype and FIFA World averaged rating.

The FWAR marginal entropy was 1.38. The FWAR conditional entropy given ACTN3 RR and ACE II genotypes was 1,27 and 1,24 respectively. However, some countries which population has high frequencies of ACTN3 RR and ACE II have low FWAR. This might be due to economic difference between countries. In general, developing countries have no ability invest a lot into childrens sport schools and national football teams what affects their FWAR. The FWAR conditional entropy given GDP has been calculated to test the hypothesis. It was 1,25 (figure 3). Finally, the correlation between GDP and genetic factors was calculated (figure 4). We found that quadratic model of ACTN3 RR genotype frequency explained 48% of log GPD per capita variance. The correlation between ACE II genotype frequency and log GPD per capita variance was less valuable (data not shown).

**Figure 3.**
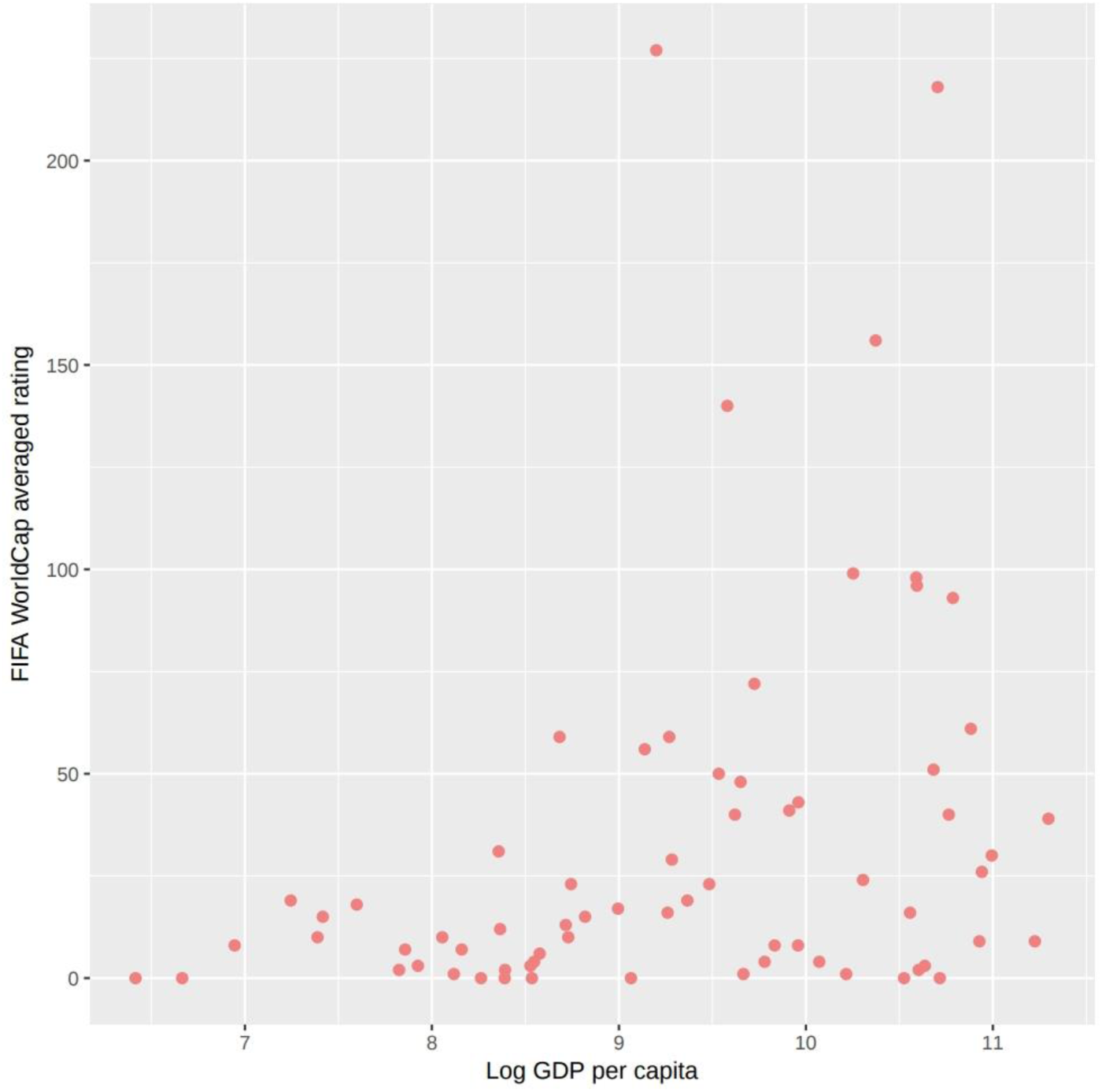
Scatter plot of correlation between log GDP per capita and FIFA World averaged rating.

**Figure 4.**
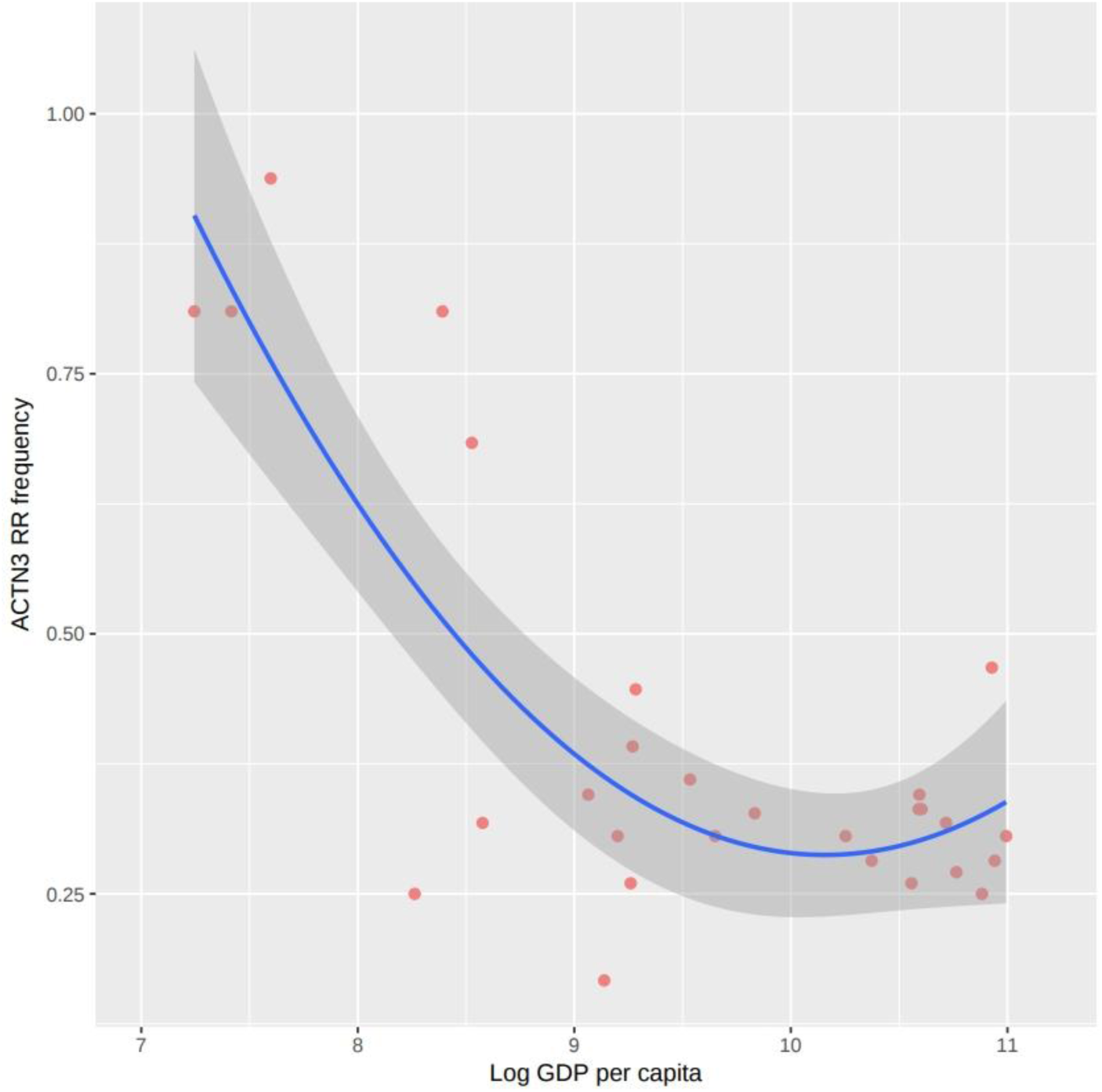
Scatter plot of correlation between log GDP per capita and ACTN3 RR genotype frequency.

## Discussion

We evaluated the contribution of ACTN3 rs1815739 and ACE indel polymorphisms on FWAR distribution entropy for the first time. Previous study showed that the distribution of ACTN3 R-allele frequencies among populations correlated with environmental conditions of their countries related to latitudinal variation, such as species richness and mean annual temperature [9]. ACE I/D genotype distribution also showed an association with longitude, with frequencies increasing eastwards and westwards from the Middle East [8]. In the present study, we found that FWAR entropy correction after adjustment on genetic factors and GDP is almost the same. Moreover, we found high correlation between ACTN3 RR genotype and GDP. It seems plausible that different factors, including GDP and genetic marker distribution, could influent on FWAR. We cannot exclude or prove any influences between these and any other unknown factors.

